# Nociceptor-enriched genes required for normal thermal nociception

**DOI:** 10.1101/053413

**Authors:** Ken Honjo, Stephanie E. Mauthner, Yu Wang, J. H. Pate Skene, W. Daniel Tracey

## Abstract

Here, we describe a targeted reverse genetic screen for thermal nociception genes of Drosophila larvae. Using laser capture microdissection and microarray analyses of nociceptive and non-nociceptive neurons we identified 275 nociceptor-enriched genes. We then tested the function of the enriched genes with nociceptor-specific RNAi and thermal nociception assays. Tissue specific RNAi targeted against 14 genes caused insensitive thermal nociception while targeting of 22 genes caused hypersensitive thermal nociception. Previously uncategorized genes were named for heat resistance (ie. *boilerman, fire dancer, oven mitt, trivet, thawb* and *bunker gear*) or heat sensitivity (*firelighter, black match, eucalyptus, primacord, jet fuel, detonator, gasoline, smoke alarm*, and *jetboil*). Insensitive nociception phenotypes were often associated with severely reduced branching of nociceptor neurites and hyperbranched dendrites were seen in two of the hypersensitive cases. Many genes that we identified were not isolated in a prior genome-wide screen, and are evolutionarily conserved in mammals.

## Introduction

The pomace fly *Drosophila melanogaster* has been developed as a robust system to study nociception (Babcock et al., 2009; Babcock et al., 2011; Im et al., 2015; Neely et al., 2010; Tracey et al., 2003). *Drosophila*, with its unparalleled genetic tools, is an excellent model to explore novel nociception genes. *Drosophila* larvae rotate along the long body axis in a corkscrew like fashion in response to noxious stimuli such as heat (>39°C) or harsh mechanical stimuli (Tracey et al., 2003). This highly stereotyped response to harmful stimuli, named nocifensive escape locomotion (NEL) or rolling, serves as a robust behavioral readout of nociception since it is specifically triggered by noxious stimuli and it is clearly distinguishable from normal locomotion and other somatosensory responses.

Several lines of evidence indicate that Class IV multidendritic (md) neurons are polymodal nociceptive sensory neurons responsible for larval thermal and mechanical nociception. The *pickpocket* and *balboa/ppk-26* genes show highly specific expression in these neurons and they are required for mechanical nociception (Gorczyca et al., 2014; Guo et al., 2014; Mauthner et al., 2014; Zhong et al., 2010). Similarly, reporter genes for specific *dTRPA1* transcripts are specifically expressed in the Class IV cells and *dTRPA1* is required for both mechanical and thermal nociception (Zhong et al., 2012). Genetic silencing of Class IV neurons severely impairs thermal and mechanical nociception behavior and optogenetic activation of these neurons is sufficient to evoke NEL (Hwang et al., 2007; Zhong et al., 2012).

## Results and Discussion

### Laser capture microdissection and microarray analyses identify 275 nociceptor-enriched genes

Genes involved in nociception are often preferentially expressed in nociceptors (Akopian et al., 1996; Caterina et al., 1997; Chen et al., 1995; Dib-Hajj et al., 1998; Mauthner et al., 2014; Nagata et al., 2005; Zhong et al., 2012; Zhong et al., 2010). Thus, to identify *Drosophila* nociceptor-enriched genes we performed laser capture microdissection to isolate RNAs of nociceptive and non-nociceptive neurons (Mauthner et al., 2014). We then performed microarray analyses on the isolated samples (Mauthner et al., 2014). We compared the gene expression profiles of nociceptive Class IV multidendritic (md) neurons to Class I md neuron profiles (Mauthner et al., 2014) as Class IV md neurons are polymodal nociceptors (their output is both necessary and sufficient for triggering larval nociception behaviors), and Class I md neurons are functionally dispensable for nociception (Hwang et al., 2007). Indeed, as internal validation of these methods, this microarray study successfully detected the enrichment of genes previously thought to be preferentially expressed in Class IV relative to Class I neurons, such as *cut, knot, Gr28b, ppk* and *balboa/ppk26* (Mauthner et al., 2014) (Table S1A).

Here, to further identify nociceptor-enriched genes, we made a side-by-side comparison of the normalized hybridization intensity between Class IV and Class I neurons for all Affymetrix probe sets, and identified 278 probe sets corresponding to 275 genes that showed a greater than two-fold higher expression in Class IV neurons in comparison to Class I neurons (Class IV / Class I > 2; p < 0.05 with Welch t-test) (Figure S1 and Table S1A).

### Nociceptor-specific RNAi screens uncover 36 genes required for larval thermal nociception

We subsequently tested the function of the nociceptor-enriched genes in thermal nociception responses. In order to test their requirement specifically in nociceptors, we used RNAi to knock down each gene in a tissue-specific pattern using the Class IV specific GAL4 driver *ppk1.9-GAL4. UAS-dicer2* was also present in the driver strain in order to enhance RNAi knockdown (Dietzl et al., 2007). A total of 419 UAS-RNAi lines were obtained from the Vienna Drosophila RNAi library (Dietzl et al., 2007), the TRiP RNAi library (Ni et al., 2009), and the National Institute of Genetics RNAi library and were used to knock down 229 of the 275 (83.3%) nociceptor-enriched genes. In control experiments, we found that the baseline nociception responses differed among the genetic backgrounds that were used to generate the different collections of UAS-RNAi strains. Thus, the progeny of our GAL4 driver strain crossed to UAS-RNAi lines from the four different collections (VDRC 1st-generation, VDRC 2nd-generation, TRiP and NIG) were each statistically analyzed in comparison to the relevant genetic controls for parental isogenic background. Progeny of each *ppk-GAL4 UAS-dicer-2* × *UAS-RNAi* cross were tested in an established larval thermal nociception assay (Tracey et al., 2003). In order to identify either insensitive or hypersensitive phenotypes, independent tests were carried out at two different probe temperatures. A 46°C probe was used to screen for insensitive phenotypes (defined as a lengthened latency to respond to the 46°C stimulus) while a 42°C probe was used to assess hypersensitivity (defined as a shortened latency to respond to this stimulus). Average latency to 46°C and 42°C thermal probe stimulation were determined for each RNAi knockdown genotype (Figure 1 and Table S1B-E). We set our initial cut-off line at the +1σ (84.13th percentile) in the insensitivity screen and -1σ (15.87th percentile) in the hypersensitivity screen, and all *ppk-GAL4 UAS-dicer-2* × *UAS-RNAi* pairs that met these cut-offs were subjected to retesting (Figure 1 and Table S1B-E). Only pairs showing significant insensitivity or hypersensitivity in comparison to the appropriate control for genetic background (p < 0.05 with Steel’s test) during the retest were considered as positive hits. Sixteen RNAi lines targeting 14 genes were identified in the insensitivity screen and 24 RNAi lines targeting 22 genes were found in the hypersensitivity screen (Table 1, Table 2, Figure 2 and S1F-M). We confirmed that these positive RNAi lines did not show the observed nociception phenotypes when crossed to *w^1118^* strain (no driver control), suggesting that the phenotypes observed in the screen were dependent on GAL4-driven expression of RNAi (Table S1N and O).

**Figure 1.**
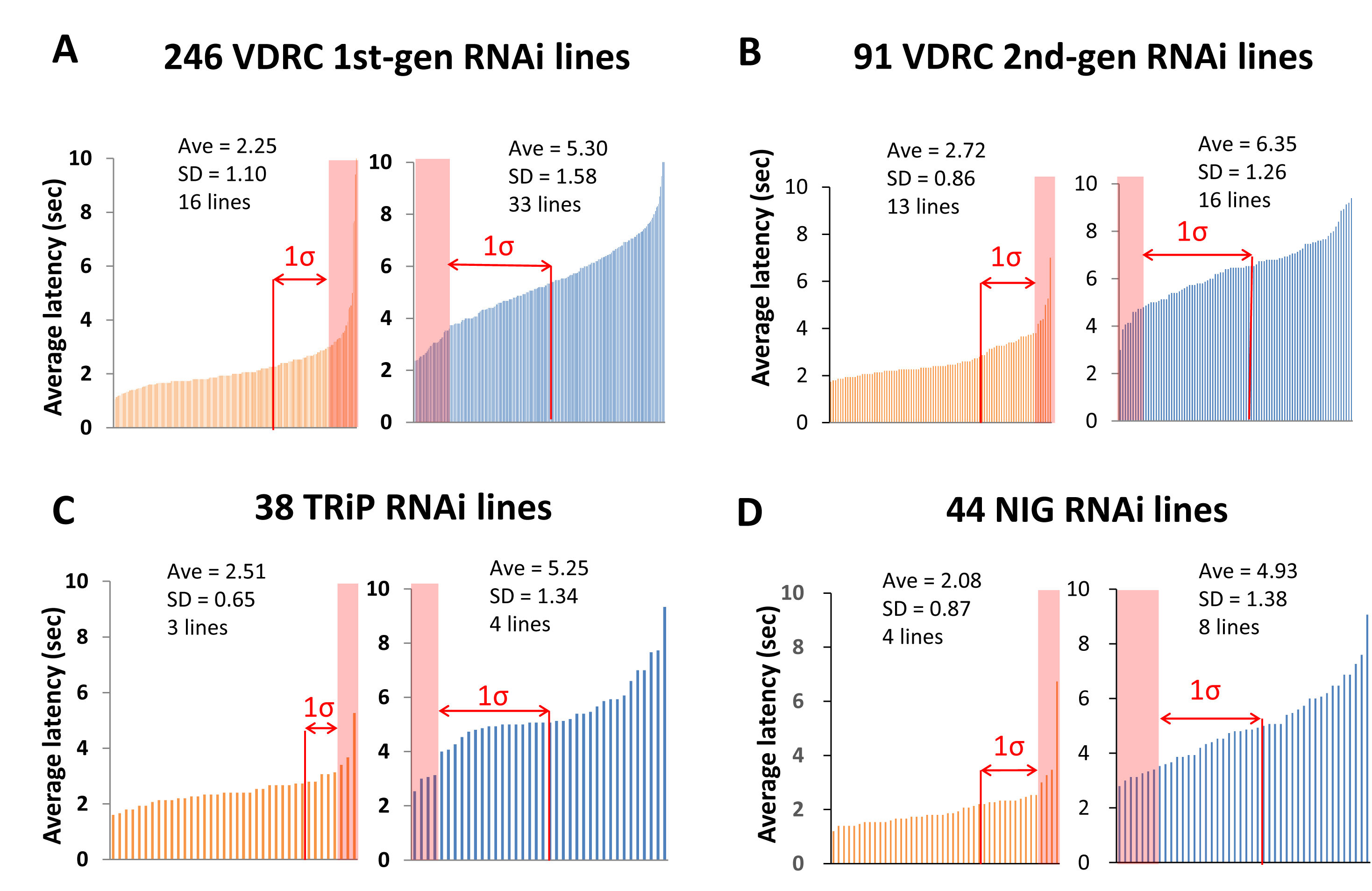
Summary of primary screen results. Summary of the insensitivity and the hypersensitivity screen with (A) 1st-generation VDRC (GD) lines, (B) 2nd-generation VDRC (KK) lines, (C) TRiP lines, (D) and NIG lines. The left chart in each panel with the orange bars show the results of the insensitivity screen with a 46oC probe. The right chart in each panel, with blue bars, shows the hypersensitivity screen results with a 42oC probe. The average latency and standard deviation of all tested lines and the number of lines which survived the initial cut-off (+1σ for the insensitivity screen and -1σ for the hypersensitivity screen) are indicated with each graph. Shaded areas indicate lines that were selected for retesting. See also Table S1B-E.

**Figure 2.**
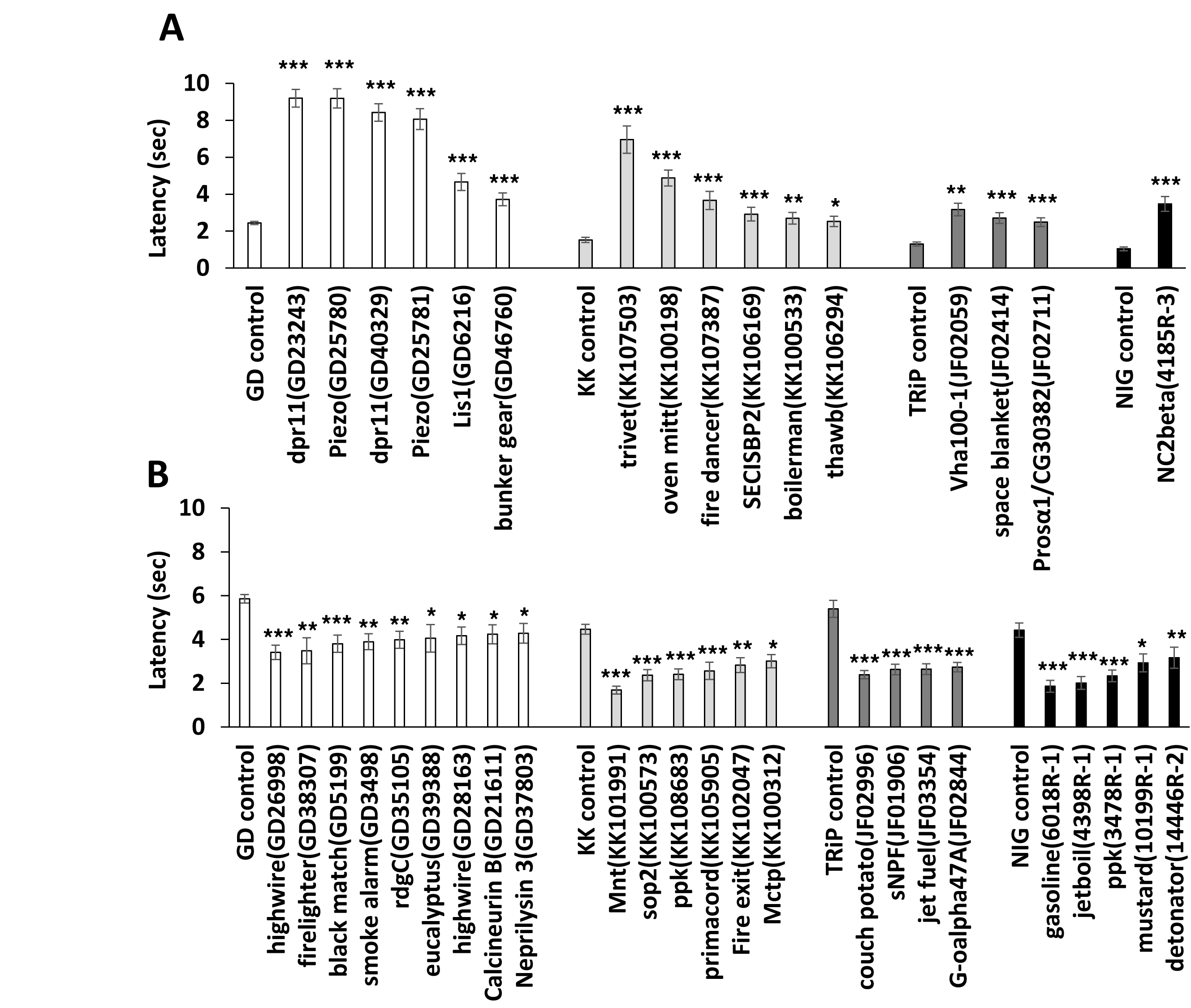
RNAi lines that showed significant insensitivity or hypersensitivity upon retesting with a larger sample size. The behavioral responses of retested lines in the hypersensitivity and insensitivity screens. Each panel shows average latency on the Y axis and the targeted genes for each genotype are listed along the × axis. (A) *ppk-*GAL4 dependent insensitive behavioral responses seen with crossing to 1st-generation VDRC (GD) lines, 2nd-generation VDRC (KK) lines, TRiP lines and NIG lines. (B) *ppk-*GAL4 dependent hypersensitive behavioral responses seen with crossing to VDRC (GD and KK) lines, TRiP lines and NIG lines. Steel’s test was used to statistically compare each genotype to its appropriate control except that Mann-Whitney’s U-test was used to perform the pair-wise comparison of NC2beta versus NIG control (n > 32; ∗ p < 0.05, ∗∗ p < 0.01 and ∗∗∗ p < 0.001). Error bars represent S.E.M. See also Table S1F-O, (Table S1 and Figure S1.

**Table 1.**
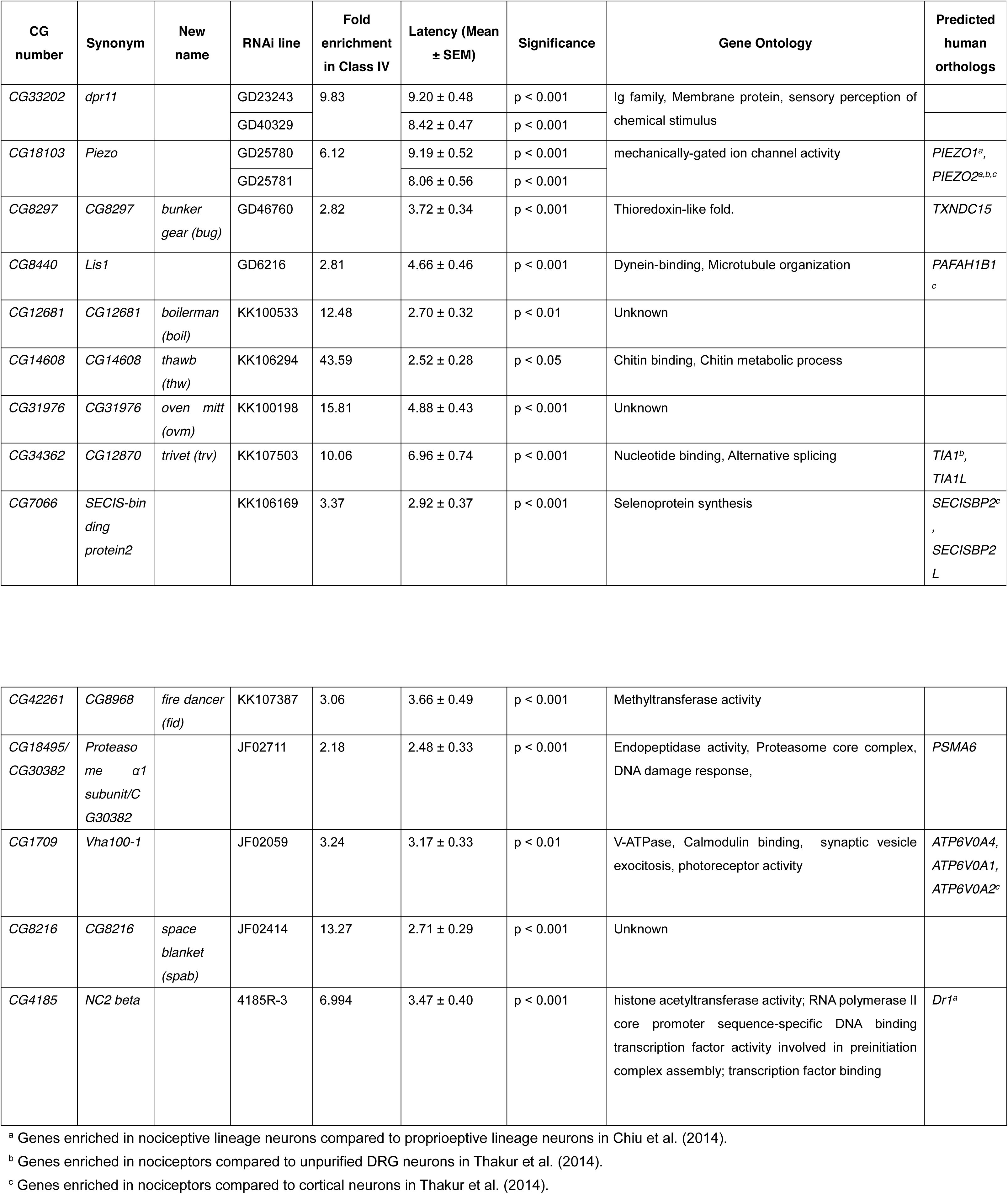
Candidate genes for insensitive nociception.

**Table 2.**
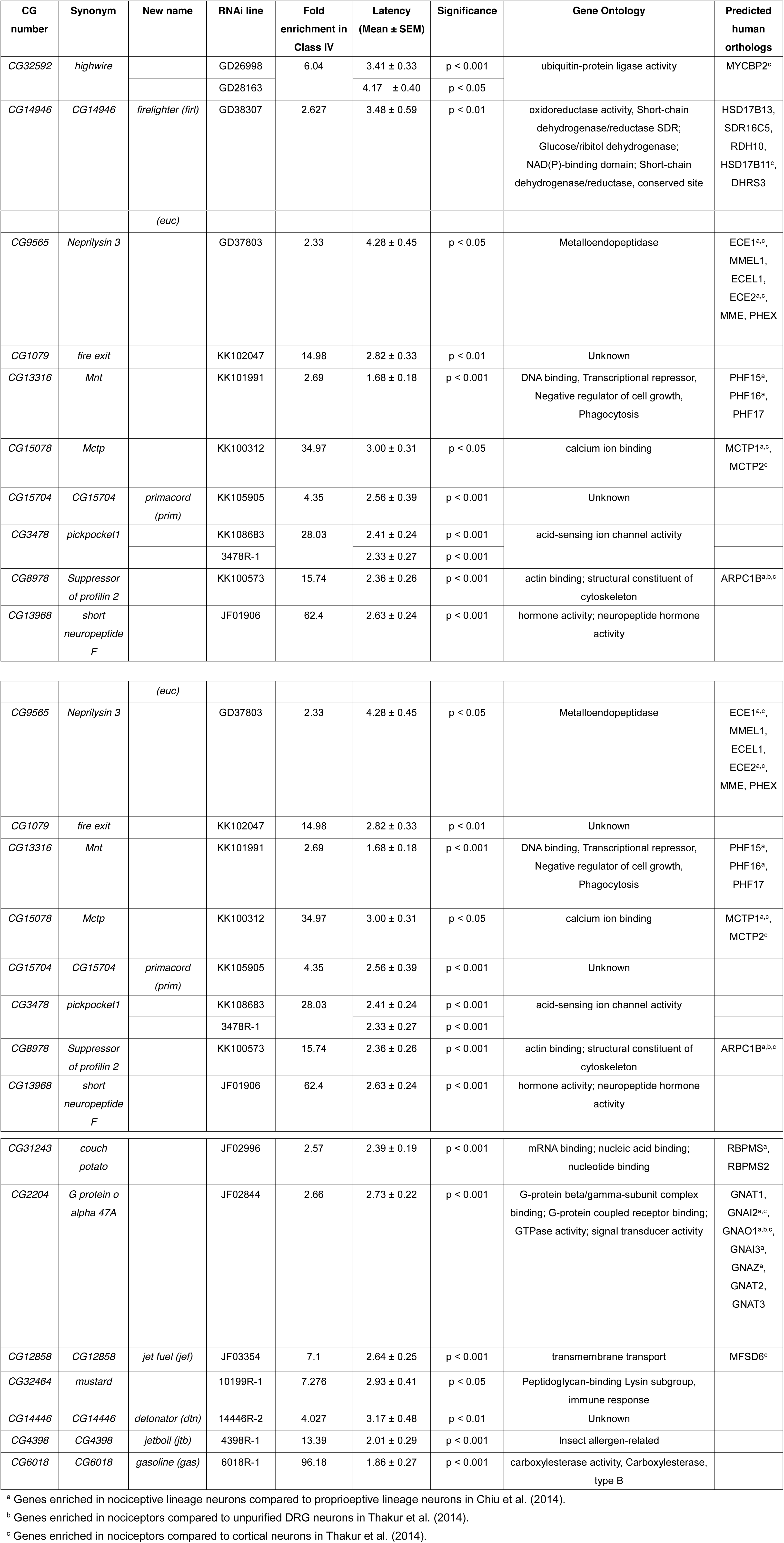
Candidate genes for hypersensitive nociception.

Neely et al. carried out a genome-wide RNAi screen for nociception genes using adult *Drosophila* (Neely et al., 2010). Neely and colleagues identified 3 genes of the 14 that we found with insensitive nociception phenotypes (*dpr11, lis-1* and *vha100-1*), and a single gene out of the 22 with hypersensitive phenotypes (*retinal degeneration C*). As this prior screen relied on thermally induced paralysis of adult flies as a surrogate for studying larval nociception behavior, it is possible that molecular mechanisms of nociception in adult flies and larval flies may distinct. In addition, Neely et al used broadly expressed knockdown approaches. It is possible that pleiotropy caused by opposing effects in distinct tissues may have masked the effects that we are able to detect with nociceptor specific knockdown. In either case, it is likely that a large number of *bona fide* nociception signaling genes remain to be discovered using larval nociception assays.

### Uncharacterized Genes Identified in the Screens

Our screen identified genes that have not been previously characterized and which remained named according to Celera Gene (CG) numbers (Table 1 and 2). Knockdown of seven CGs caused insensitivity and nine CGs caused hypersensitivity. Thus, given these loss of function phenotypes we have named each of the heat-insensitivity screen genes (*boilerman* (*boil*), *fire dancer* (*fid*)), *oven mitt* (*ovm*) and *trivet* (*trv*), *thawb* (*thw*), *bunker gear* (*bug*), *space blanket* (*spab*)) (Table 1). And we have also named uncharacterized genes that were identified in the hypersensitivity screen (*firelighter* (*firl*), *black match* (*bma*), *eucalyptus* (*euc*), *primacord (prim), jet fuel (jef), detonator (dtn)* and *gasoline (gas)*), *smoke alarm (smal)*), *jetboil (jtb)*) (Table 2).

### Genes required for Class IV neuron morphogenesis

*Lis1*, which showed an insensitive nociception phenotype in our screen (Figure 2A and Table 1), is a component of a dynein-dependent motor complex that is known to play a role in dendrite and axonal morphogenesis in Class IV neurons (Satoh et al., 2008). Similarly, a reduced dendrite phenotype in another study was associated with insensitive nociception behaviors (Stewart et al., 2012). These observations raised the possibility that nociception phenotypes detected in our screen might also be associated with defects in dendrite morphogenesis.

To test for this possibility, we used confocal microscopy to observe and quantify the dendritic coverage of CD8-GFP expressing Class IV neurons in all of the genotypes that showed insensitive or hypersensitive thermal nociception phenotypes (Figure 3 and 4). Consistent with the previous study, Lis1 RNAi showed severe defects in dendrite morphogenesis (Figure 3). Significantly reduced nociceptor dendrites were also found with RNAi lines targeting *piezo, oven mitt, trivet, fire dancer, SECISBP2, pros-alpha1* and *NC2beta* (Figure 3). Among hypersensitive hits, *smoke alarm* and *G-oalpha47A* RNAi resulted in significantly increased dendrite phenotypes, which potentially contributes to their hypersensitive nociception phenotypes (Figure 4).

**Figure 3.**
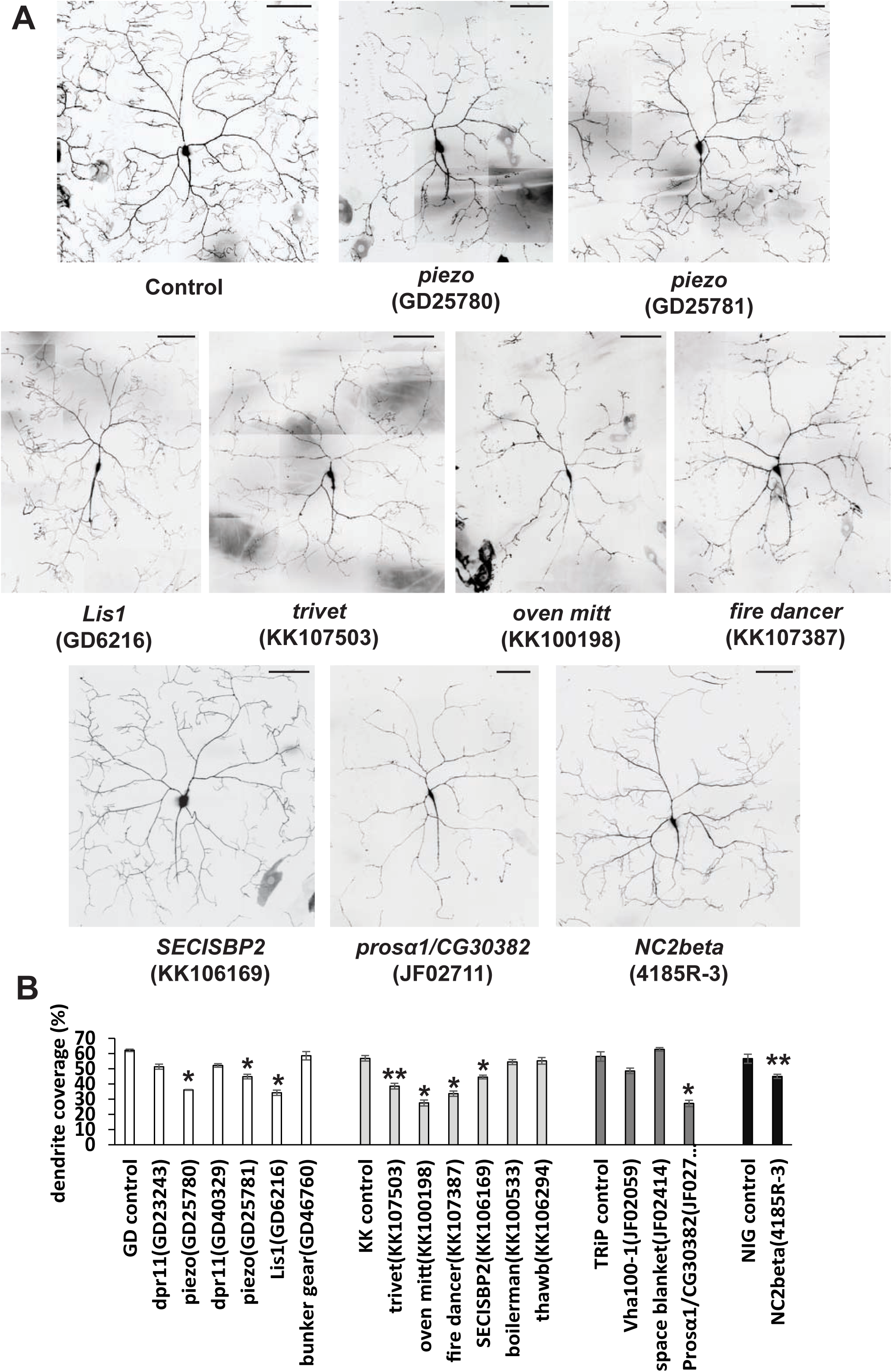
Genes targeted with RNAi that showed a reduced dendrite phenotype. (A) Representative composite images of the dendritic structure of Class IV ddaC neurons in thermal nociception insensitive RNAi lines that also showed significantly reduced dendritic coverage. Scale bars represent 100 μm. (B) Quantification of dendritic coverage for insensitive RNAi lines. Steel’s test was used for statistical comparisons between each genotype and controls, except that Mann-Whitney’s U-test was used to compare NC2beta and NIG control (n > 4; ∗ p < 0.05 and ∗∗ p < 0.01). Error bars represent S.E.M. For representative images of RNAi lines that did not show significantly reduced dendrite, see Figure S2.

**Figure 4.**
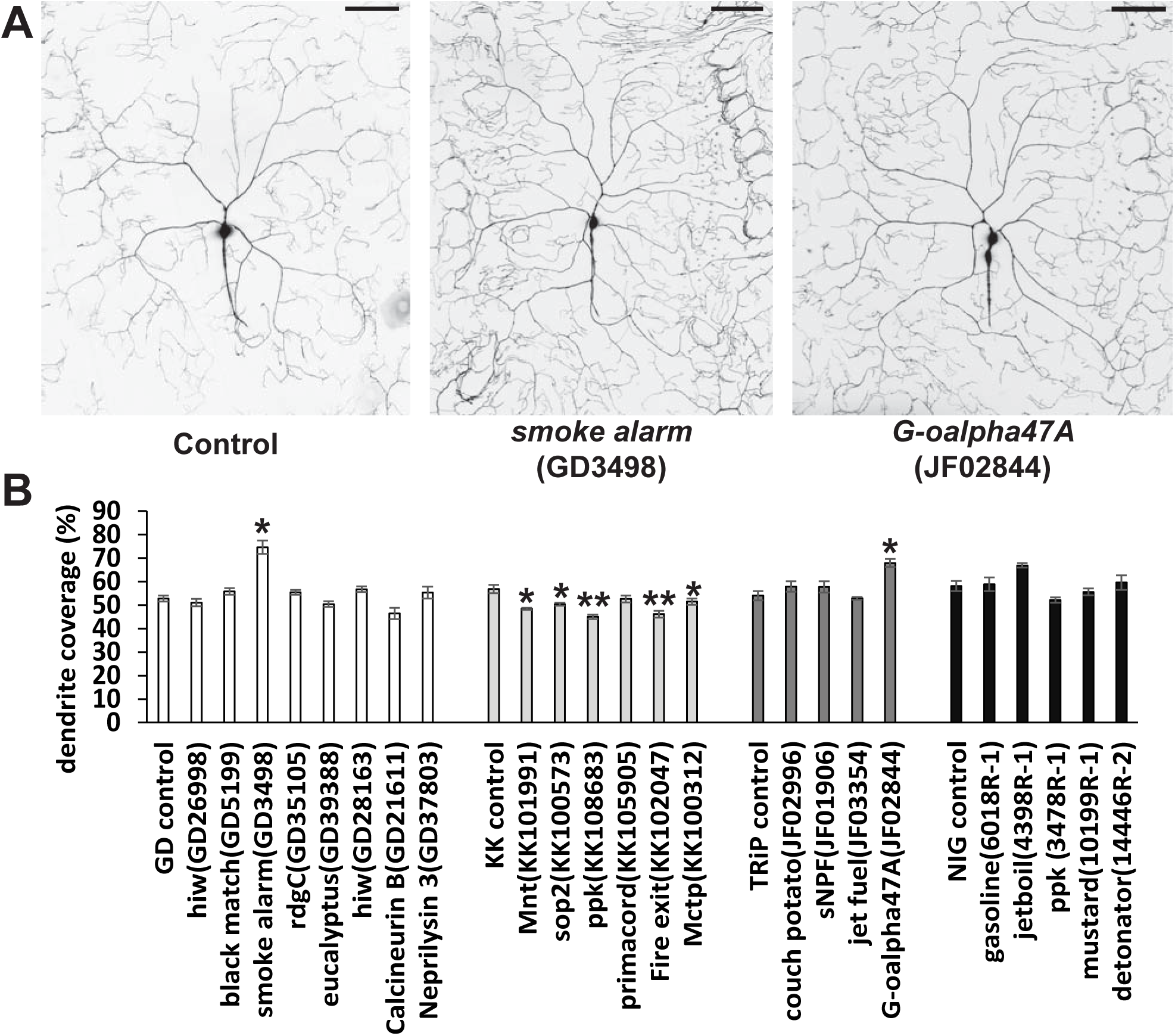
Genes targeted with RNAi that showed a an increased dendrite coverage phenotype. (A) Representative composite images of the dendritic field of Class IV ddaC neurons in RNAi lines with hypersensitive thermal nociception animals that also showed significantly increased dendritic coverage. Scale bars represent 100 μm. (B) Quantification of dendritic coverage for hypersensitive candidate RNAi lines. Steel’s test was used for statistical comparisons between each genotype and controls, except that Mann-Whitney’s U-test was used to compare NC2beta and NIG control (n > 4; ∗ p < 0.05 and ∗∗ p < 0.01). Error bars represent S.E.M. For representative images of RNAi lines that did not show significantly increased dendrite coverage, see Figure S3. The effects seen in the KK collection was not a consequence of unintended Tio expression (See supplemental experimental procedures, (Table S2 and Figure S2)

In contrast, some hypersensitive hits actually showed a mild (but statistically significant) reduction in dendrite coverage (*Mnt, sop2, ppk, fire exit* and *Mctp*) (Figure 4). Thus, perhaps not surprisingly, the degree of dendrite branching cannot be perfectly correlated with noxious heat sensitivity. An interesting possibility is that targeting these genes with RNAi results in hypersensitivity due to effects in another compartment of the cell such as axons and/or synapses. Or alternatively, hypersensitivity in the dendrites masks a pleiotropic morphologically reduced dendrite phenotype.

*highwire*, one of hypersensitive candidates (Figure 2B and Table 2), has been shown to be important for dendrite and axonal morphogenesis of Class IV neurons (Wang et al., 2013). Interestingly *hiw* RNAi did not cause reduced dendritic arbors that have been reported with strong loss of function alleles for *hiw.* Our detailed analyses of *hiw* indicate that the hypersensitive nociception phenotype is more sensitive to *hiw* dosage than are the previously reported dendrite phenotypes (Honjo and Tracey, unpublished observations). As in the case of *hiw*, RNAi knockdown often results in phenotypes that are less severe than those that would be observed with null alleles.

Indeed, there are other caveats to be considered when using an RNAi screening methodology. The incomplete knockdown effect can also result in false negatives, which are estimated to occur in up to 40% of the UAS-RNAi strains in the major collection of strains used in our screen. Thus, the lack of a phenotype in our screen cannot be used to conclusively infer a lack of function for a particular gene of interest. As well, false positives may occur, presumably due to off target effects. When the UAS-RNAi used in conjunction with *UAS-dicer-2* (as in our experiments) the effectiveness of knockdown is enhanced, and off-target effects are seen in approximately 6% of lines (when tested in the very sensitive crystalline lattice of the eye, or in the notum). Applying this estimate to the 36 genes implicated by our screen cautions that two or more of the candidates may represent false positives.

### Nociception genes are evolutionarily conserved in mammals

Twenty out of the thirty-six of the genes that are implicated here in nociception have clearly predicted mammalian orthologues (Table 1 and 2). Interestingly, published evidence supports roles for some of these orthologues in regulating mammalian nociception. Nociceptor and thermoreceptor-specific knock-out of *MYCBP2*, a mammalian homologue of *highwire*, shows prolonged thermal hypersensitivity with formalin-induced hyperalgesia (Holland et al., 2011). Knockdown of *highwire* caused an intriguingly similar hypersensitivity to heat (Figure 2B and Table 2).

RNAi targeting *Neprilysin-3*, encoding a neutral endopeptidase, showed hypersensitive nociception (Figure 2B and Table 2). Loss of ECE2, one of six predicted mammalian homologues of Neprilysin-3, has been also implicated in hypersensitive nociception as a knock-out mouse for ECE2 exhibits thermal hypersensitivity and rapid tolerance to morphine induced analgesia (Miller et al., 2011). In addition, knock-out of MME (aka NEP), another homologue of Neprilysin-3, causes thermal hyperalgesia (Fischer et al., 2002). These results thus raise the possibility that inhibitory nociceptive functions of Neprilysin-3 may be conserved between flies and mammals.

To our knowledge, the remaining conserved genes that our studies implicate in nociception have not been functionally implicated in mammalian pathways. However, it is very intriguing that many of these conserved candidate genes are more highly expressed in nociceptors compared to non-nociceptive neurons or other tissues (Chiu et al., 2014; Goswami et al., 2014; Thakur et al., 2014). The orthologues of the seventeen out of twenty candidate nociception genes that came out from our study have been reported to show significantly enriched expression in nociceptive sensory neurons (Table 1 and 2). These genes will thus be particularly promising targets to identify previously uncharacterized molecular pathways involved in nociception.

Two ion channel genes from our screen have been implicated in mechanical nociception, *ppk* and *Piezo* (Figure 2, Table 1 and 2). Since studies of *Piezo* in Class IV neurons have observed defective phenotypes in mechanical nociception assays but not in thermal nociception assays (Kim et al., 2012) it is surprising that *Piezo* RNAi causes thermal insensitivity. The apparent discrepancy may be because the two *Piezo* RNAi strains used in this study specifically target low-abundance exons that were previously annotated as an independent gene, *fos28F* (Graveley et al., 2011). And in our microarray dataset we found enriched expression for “*fos28F”* in Class IV neurons but not for *piezo.* Our interpretation of these microarray data is that a nociceptor specific transcript for *piezo* exists and it contains sequences from the previously annotated gene *fos28F.* Both of the RNAi constructs that target the *fos28F/piezo* exons caused gross abnormalities in dendritic and axonal gross morphology (Figure 3 and data not shown). RNAi lines targeting canonical *piezo* exons do not cause a similar thermal nociception phenotype (KH unpublished observations). Thus, the nociceptor-specific knockdown of the low-abundance transcriptional variant containing exons from *fos28F* appears to disrupt the morphology and thermal nociception capacity of Class IV neurons.

It was also unexpected that two independent *ppk* RNAi strains collected from different libraries showed hypersensitive thermal nociception phenotypes. This contrasts with the severely insensitive mechanical nociception phenotypes that occur with the loss of *ppk* (Zhong et al., 2010). Our previous studies did not detect thermal hypersensitivity due to the testing with a single higher probe temperature (46°C). The new finding that *ppk* RNAi causes thermal hypersensitivity highlights the importance of using the 42°C probe temperature in the search for hypersensitive phenotypes. In addition, the results indicate that modality specific phenotypes within the larval nociceptors can be of opposite sign. It is interesting to note that *ppk* mutants have been found to show a locomotion phenotype in which the animals crawl rapidly in a straight line across the substrate with very infrequent turning (Ainsley et al., 2003). A similar form of locomotion is also seen in a second phase of nociception behavior that immediately follows rolling behavior (NEL) (Ohyama et al., 2013). Thus, it is interesting to speculate that the locomotion phenotype of *ppk* mutants is actually a consequence of a hypersensitive process in the nociceptors. This in turn may be causing the larvae to continuously manifest the fast crawling phase of nociception escape.

In conclusion, we have carried out a large-scale screen that combines molecular approaches to identify cell type enriched nociceptor RNA with *in vivo* functional studies of the same identified RNAs in a phenotypic screen. This approach has led to the identification of a set of nociception genes. Many of these genes are evolutionarily conserved and also show enriched expression in mammalian nociceptors, future studies will reveal the physiological importance and molecular mechanisms that depend on these molecules.

## Experimental procedures

### Laser Capture Microdissection and Microarray analysis

Detailed methods for our Laser Capture Microdissection and Microarray analysis were previously described in Mauthner et al. (Mauthner et al., 2014).

### Thermal nociception screen

Three males of each RNAi strain were crossed to six virgin females of *ppk1.9-GAL4; UAS-dicer2* strain in a standard molasses cornmeal food vial, and incubated for 5 to 7 days at 25°C prior to harvest of the F1 wandering third instar larvae. Control crosses (a control strain crossed to ppk1.9-GAL4; UAS-dicer2) were performed side-by-side.

Thermal nociception assays were performed as described previously (Hwang et al., 2012; Tracey et al., 2003; Zhong et al., 2012). To detect insensitivity and hypersensitivity phenotype efficiently, each *UAS-RNAi* × *ppk1.9-GAL4; UAS-dicer2* pair was tested by using two different probe temperatures: A custom-made thermal probe heated to 46°C was used to test insensitivity and the probe heated to 42°C was used for hypersensitivity.

### Quantifying dendrite coverage of Class IV neurons

Dendritic field coverage was quantified on composite images of maximum intensity projections of confocal micrographs and quantified as described previously (Stewart et al., 2012) with slight modifications. Images of ddaC neurons were overlaid with a grid of 32 × 32 pixel squares (14 × 14 μm), and squares containing dendritic branches were counted to calculate a dendritic field coverage score (i.e. the percentage of squares containing dendritic branches / the total number of squares). Counting dendrite-positive squares was done with a Matlab custom code and then manually curated to eliminate false-positives and false-negatives. One or two neurons from each imaged animal were analyzed.

### Statistical analyses

All pair wise comparisons were performed with Mann-Whitney’s U-test. For multiple comparisons, Steel’s test (non-parametric equivalent of Dunnet’s test) was used. Statistical analyses were performed in the R software and Kyplot.

## Accession numbers

The ArrayExpress (https://www.ebi.ac.uk/arrayexpress/) accession number for the microarray dataset reported in this study is E-MTAB-3863.

## Acknowledgements

We thank the Bloomington *Drosophila* Stock Center (NIH P40OD018537), the Vienna *Drosophila* RNAi Stock Center, TRiP at Harvard Medical School (NIH/NIGMS R01-GM084947), and the National Institute of Genetics Fly Stock Center. We are grateful to Dr. Kyriacou for *UAS-tio* strain. Eashan Kumar and Stephen Jeffirs assisted with crosses. This work was supported by grants from the National Institutes of Health (U24NS51870 to J.H.P.S. and R01GM086458 to W.D.T.) and JSPS KAKENHI (26890025 to K.H.). K.H. was supported by fellowships from the Uehara Memorial Foundation, the Ruth K. Broad Biomedical Research Foundation and the Japan Society for the Promotion of Science.

### Author Contributions

KH performed experiments, analyzed data and wrote the manuscript, SEM performed experiments, analyzed data and wrote the manuscript, YW performed LCM experiments, JHPS directed the LCM core for the NIH Microarray Consortium, WDT conceived the project, analyzed data, and wrote the manuscript.

## Supplemental information

### Supplemental legend

**Table S1. Identified nociceptor-enriched genes and screen result (related to Figure 1 and 2).**

(A) A list of nociceptor-enriched genes. CG number, enrichment in Class IV (Class IV/Class I), p-value with Welch’s t-test and synonyms are shown for each gene. (B-E) Initial screen results. VDRC 1st-gen (GD) RNAi lines (B), VDRC 2nd-gen (KK) RNAi lines (C), TRiP RNAi lines (D) and NIG RNAi lines (E). The ID of the RNAi line in each library, and CG number, annotated name, Class IV enrichment of the target gene are included. Average latency, SEM, and the number of larvae tested are separately shown for the insensitivity screen (46 °C) and hypersensitivity screen (42 °C). RNAi lines that fell above the cut-off line for the insensitivity screen are highlighted in orange and RNAi lines that fell below the cut-off line in hypersensitivity screen are highlighted in blue. (F-M) Retest results. Retest results of the insensitive hits from VDRC 1st-gen (GD) RNAi lines (F), hypersensitive hits from VDRC 1st-gen (GD) RNAi lines (G), insensitive hits from VDRC 2nd-gen (KK) RNAi lines (H), hypersensitive hits from VDRC 2nd-gen (KK) RNAi lines (I), insensitive hits from TRiP RNAi lines (J), hypersensitive hits from TRiP RNAi lines (K), insensitive hits from NIG RNAi lines (L) and hypersensitive hits from NIG RNAi lines (M). Lines are listed from strongest to weakest phenotypes in the initial screen. The number of larvae tested, average latency, SEM and p-value when compared to a control score in the retest are shown. The ID of the RNAi line in each library, and CG number, annotated name, Class IV enrichment of the target gene and average latency in the initial screen are also included. RNAi lines whose insensitive and hypersensitive phenotype were retested with statistical significance are highlighted in orange and blue, respectively. (N and O) Results of no driver controls. Positive hit RNAi lines were crossed to *w^1118^* strain and tested for the phenotype that was observed in retests when combined with *ppk-GAL4 UAS-dicer2* strain. The number of larvae tested, average latency, SEM and p-value when compared to a control score in the retest are shown.

**Table S2.**
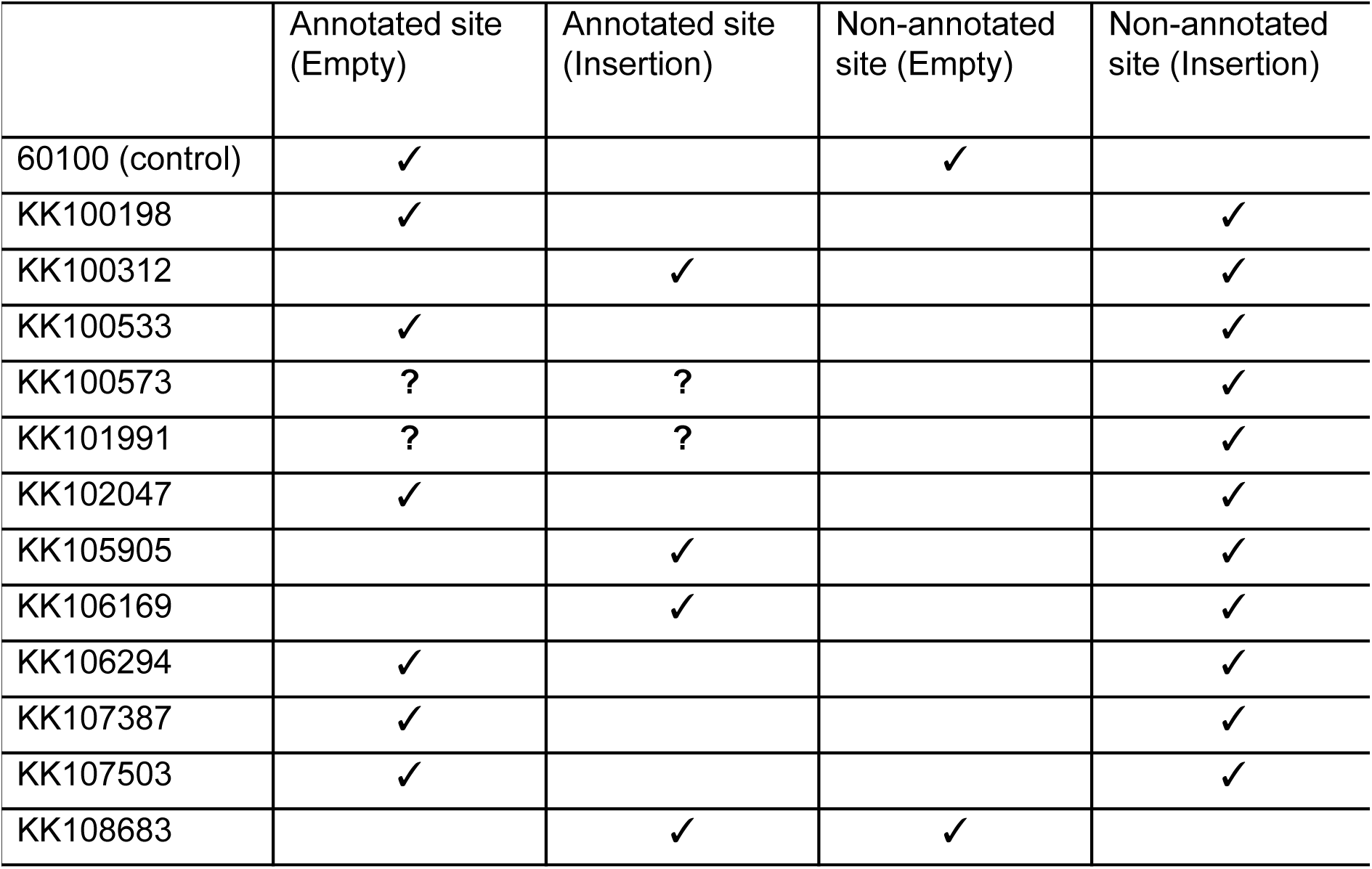
PCR verification of insertion sites for our KK line hits (related to Figure 2) PCR reactions were carried out to determine the presence or absence of UAS-RNAi insertions at the annotated attP site or the “non-annotated site” according to the methods described in Green et al (2014). Check marks indicate a positive PCR result confirming either the presence of an insertion or an empty attP site. The PCR results were not informative for two of the lines (KK100573 and KK101991) at the annotated site as neither reaction for this site produced a positive PCR result. The uncertainty for the annotated site in these lines are indicated as question marks. Note that insertions at the annotated site have the potential to be problematic due to unintended over-expression of *tio.*

**Figure S1.**
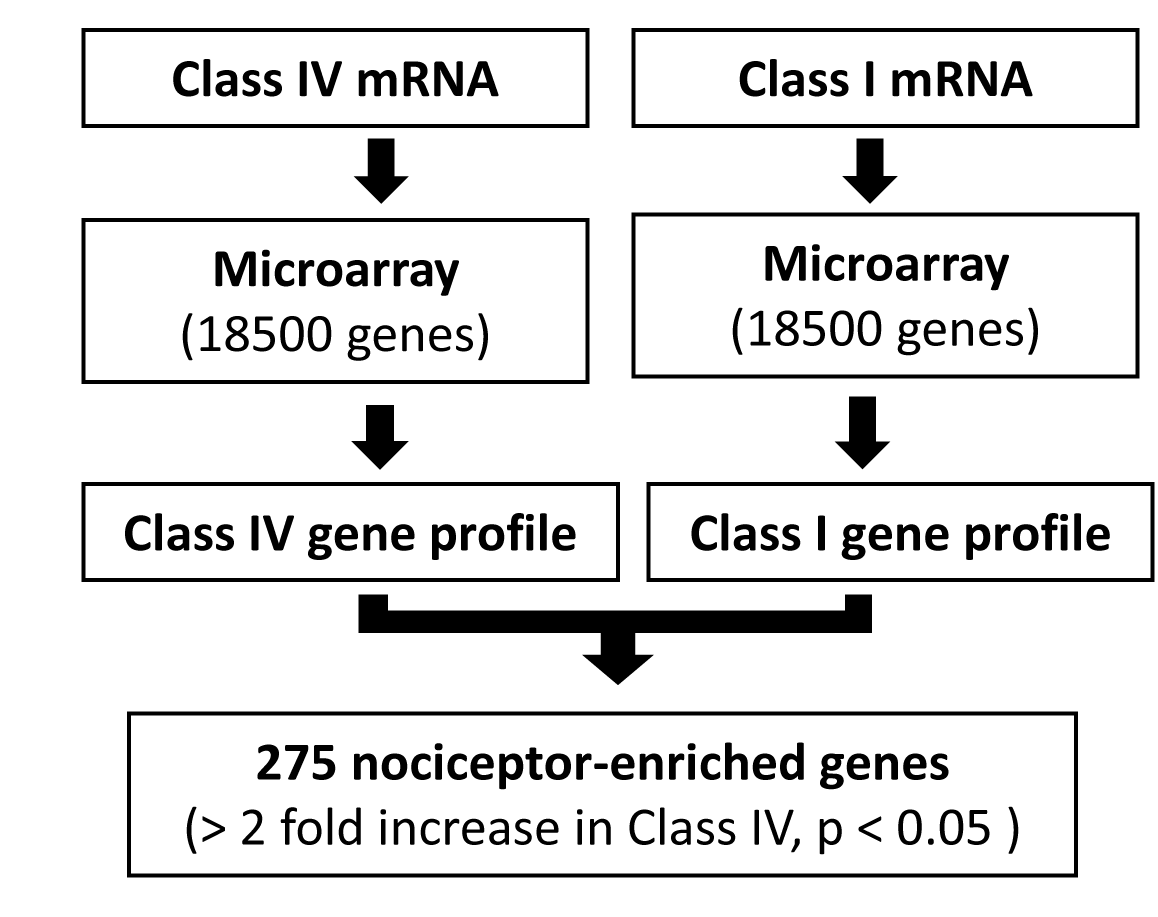
Identification of Class IV enriched genes (related to Figure 1) A flow chart of the comparative microarray analysis to identify nociceptor-enriched genes. See also Figure1 and Table S1A.

**Figure S2.**
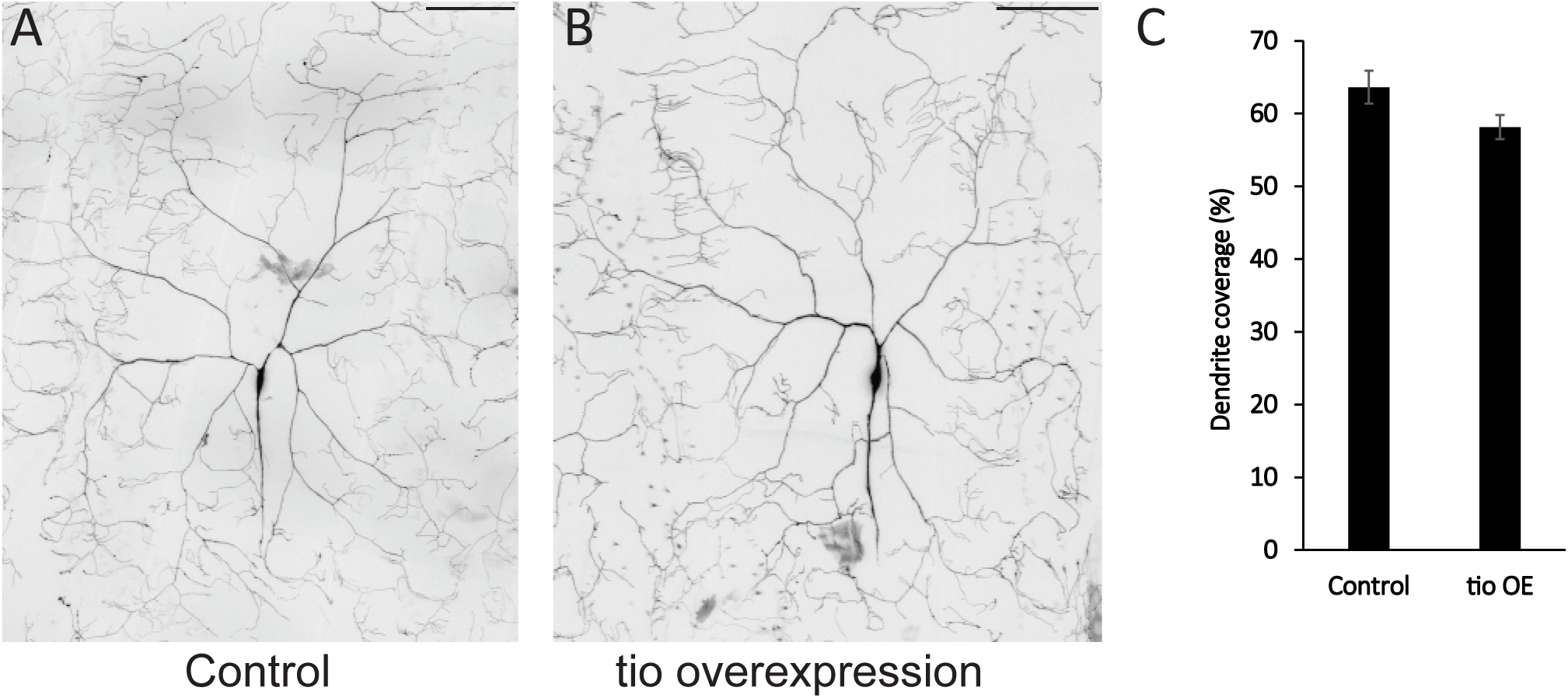
Dendritic morphology of Class IV neurons overexpressing tio gene (related to Figure 4) (A and B) Representative images of the dendritic structure of ddaC Class IV neurons. Control *(ppk-GAL4 UAS-mCD8::GFP; UAS-dicer2 × w^1118^)* and tio overexpression *(ppk-GAL4 UAS-mCD8::GFP; UAS-dicer2 × UAS-tio)*. Scale bars represent 100 μm. (C) Quantified dendrite coverage of ddaC Class IV neurons overexpressing tio. No statistical difference was detected in conmarison to control neurons (p > 0.1, n = 7 and 8, Mann-Whitney’s U-test). Error bars represent SEM.

**Figure S3.**
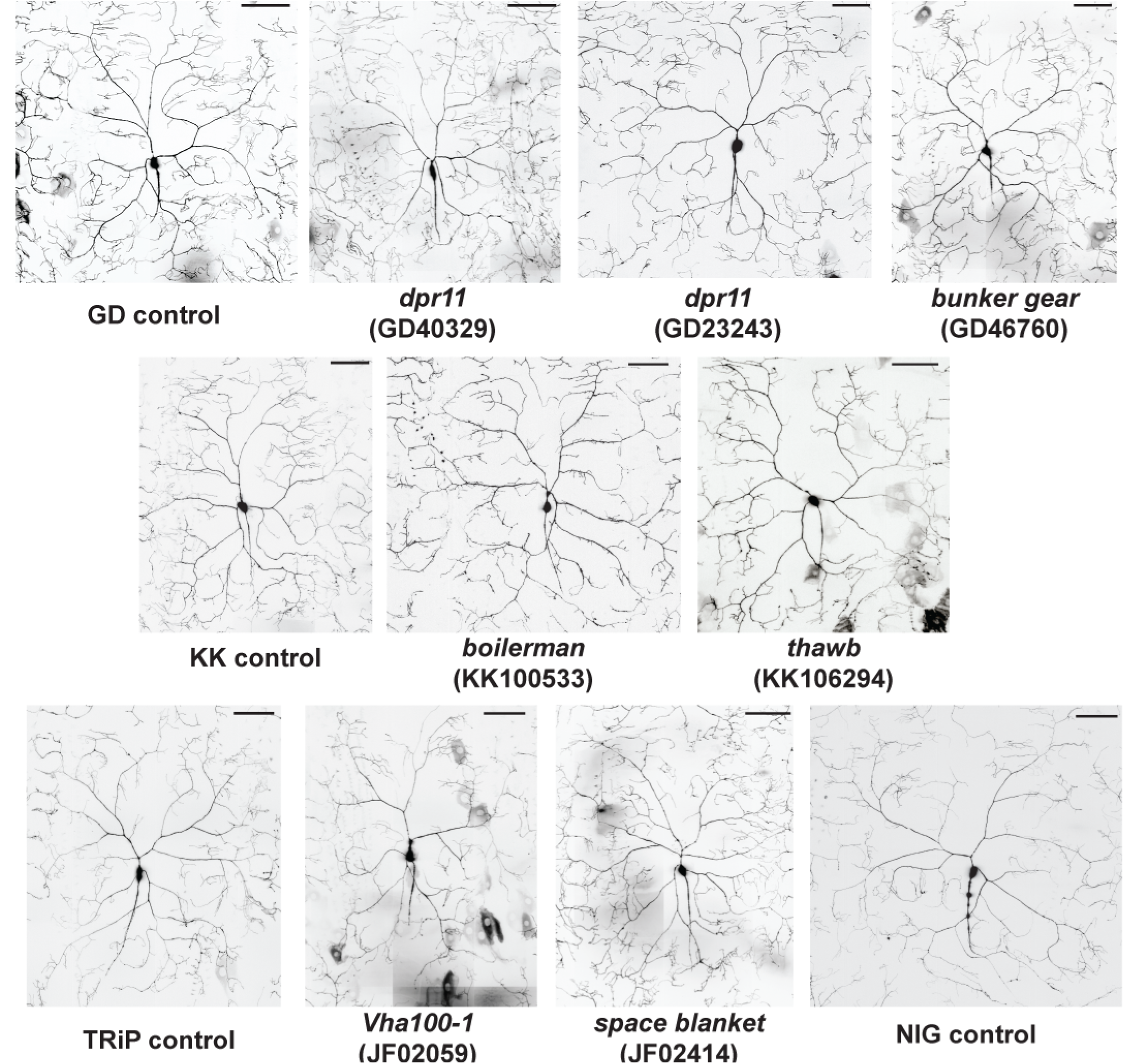
RNAi strains that showed insensitive thermal nociception but unaltered dendritic morphology of Class IV neurons (related to Figure 3) Representative pictures of the dendritic structure of ddaC Class IV neurons in RNAi animals that exhibited insensitive thermal nociception in our screen and control animals. See Figure 3 for quantified data. Scale bars represent 100 μm.

**Figure S4.**
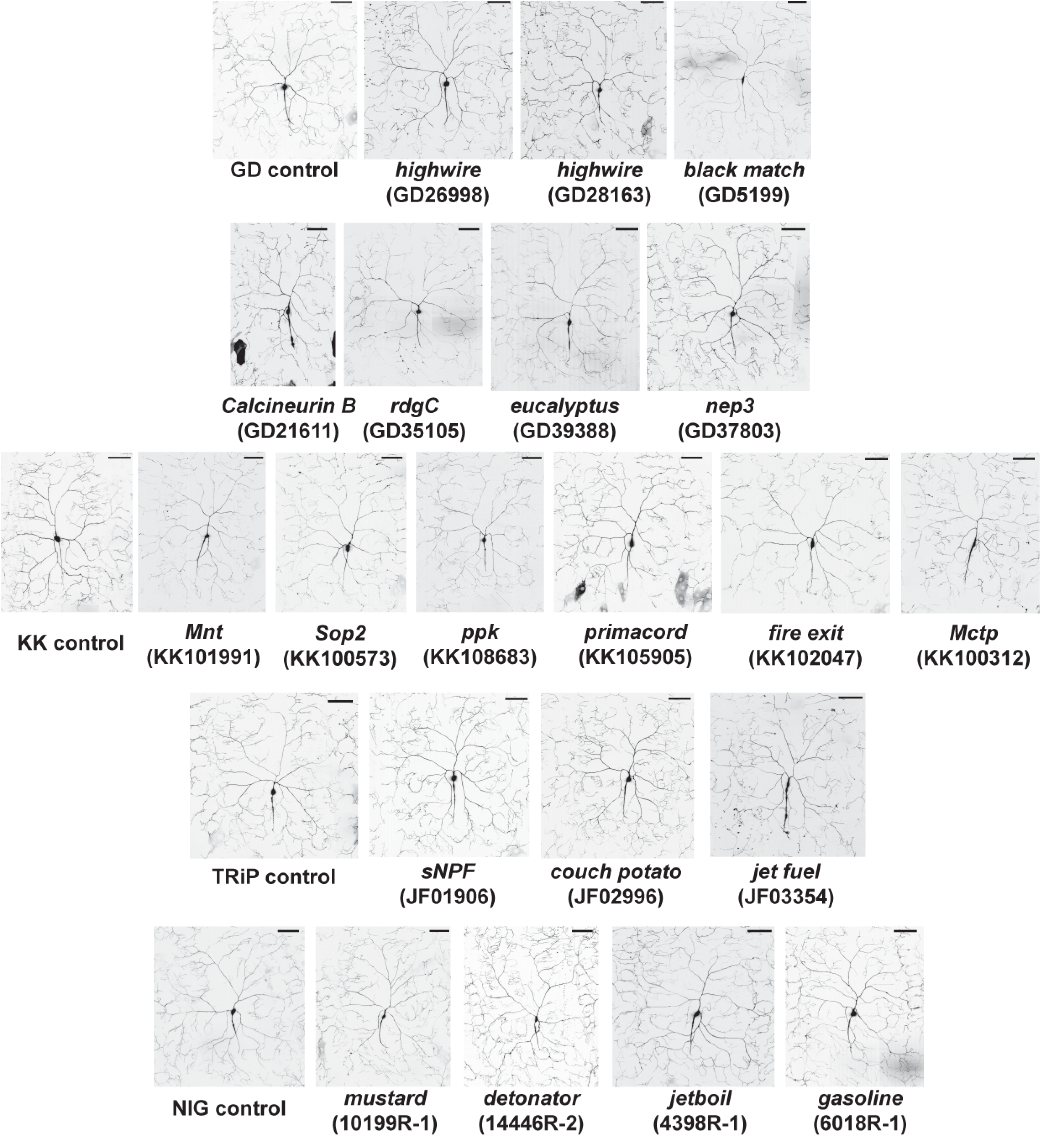
RNAi strains that showed hypersensitive thermal nociception but unaltered dendritic morphology or mild reduction in coverage of Class IV neurons (related to Figure 4) Representative pictures of the dendritic structure of ddaC Class IV neurons in RNAi animals that showed hypersensitive thermal nociception in our screen and control animals. See Figure 4 for quantified data. Scale bars represent 100 μm.

## Supplemental experimental procedures

### Fly strains

All UAS-RNAi lines tested in the nociceptor-specific RNAi screen are listed in Table S1B-E. The VDRC provides a computational prediction for the number of possible off-target effects for each line in the collection. As a precaution against off-target effects, we did not include any line with greater than two potential off-targets in our genetic screen. VDRC *isow* line, VDRC 60100, *yv; attP2* and *w^1118^* strains crossed to *w; ppk1.9-GAL4; UAS-dicer2* strain were used as controls for VDRC 1st-gen RNAi (GD) lines, VDRC 2nd-gen RNAi (KK) lines, TRiP RNAi lines and NIG RNAi lines, respectively. *ppk1.9-GAL4 UAS-mCD8::GFP; UAS-dicer2* was used for dendrite imaging.

### Behavioral experiments

Different sets of larvae were used for 46°C and 42°C tests. In the initial screen, at least 15 larvae were tested for each *UAS-RNAi* × *ppk1.9-GAL4; UAS-dicer2* pair. Average latency to respond to the thermal probe stimulation was calculated and compared to the latency of control crosses. Crossed progeny from driver to RNAi strains that showed significantly longer latency to respond to 46°C probe or shorter latency to 42°C probe than controls were retested. Approximately 45 larvae were tested in the repeated testing round. The latency data for the second round of testing for the progeny of each *UAS-RNAi × ppk1.9-GAL4; UAS-dicer2* cross was compared to pooled latency data of control crosses that were tested side-by-side with the RNAi crosses, and RNAi strains whose phenotype held up were determined as positive hits. Steel’s test (non-parametric equivalent of Dunnet’s test) was used for the statistical comparisons, except that Mann-Whitney’s U-test was used to perform the pair-wise comparison between controls and NC2beta RNAi shown in Figure 2A and 3B.

### Dendrite imaging

Each RNAi line was crossed to *ppk1.9-GAL4 UAS-mCD8::GFP; UAS-dicer2*. Wandering third instar larvae were harvested and anesthetized by submersion in a drop of glycerol in a chamber that contained a cotton ball soaked by a few drops of ether. Class IV neurons in the dorsal cluster (ddaC neurons) in segments A4-6 were imaged on Zeiss LSM 5 Live with a 40x/1.3 Plan-Neofluar oil immersion objective. A series of tiled Z-stack images were captured and assembled by the Zeiss software package to reconstruct the entire dendritic field of ddaC neurons. Maximum intensity projections were then generated from Z-stack images.

### Testing the effects of *tiptop*

It has been recently reported that VDRC 2nd-generation RNAi strains have an unexpected confound (Green et al., 2014) in which some insertions cause unintended overexpression of the *tiptop* (*tio*) gene. To test for this confounding effect, we performed a PCR validation for positive hits from the VDRC 2nd-generation library and found that only one insensitive candidate line (KK106169) and three hypersensitive candidate lines (KK100312, KK108683 and KK105905) possess the transgene integration at the annotated site that is predicted to cause overexpression of *tio* gene (Table S2). Two other hypersensitive candidates showed inconclusive PCR results, which might be accompanied by an integration and/or rearrangement of the annotated integration site (Table S2). If *tio* overexpression on its own were to cause a non-specific nociception phenotype we would expect to observe defective nociception phenotypes in a higher fraction of lines from the KK collection. Thus, the nociception phenotypes in majority of our hits from the KK collection cannot be explained by unintended *tio* expression. PCR for detection of KK lines that may affect expression of *tiptop* were performed as previously described (Green et al., 2014). In addition, we found no effect of overexpressing of tio on dendrite morphogenesis (Figure S2).

